# Attention, Emotion, and Authenticity: Eye-Tracking Evidence from AI vs. Human Visual Design

**DOI:** 10.1101/2025.10.22.684027

**Authors:** Murat Aytaş, Kamil Kerem Yıldırım, Sefer Kalaman, Mustafa Böyük, Esma Nur Cerinan Otovic

## Abstract

This study investigates how viewers perceive, attend to, and emotionally respond to AI-generated versus human-created visual content, integrating multimodal data from eye-tracking, facial-coding, and self-report surveys. The sample consisted of 136 undergraduate and graduate students enrolled in a graphic design program at a public university. Participants viewed a series of static and video stimuli produced either by human designers or artificial intelligence systems. Gaze behavior (fixation count, duration, and saccade length), emotional reliability (k-coefficient from RealEye facial-coding), and attitudinal evaluations were analyzed through both parametric and nonparametric statistical tests. The results reveal that human-made visuals elicited longer viewing durations (M = 7035 ms), higher fixation counts (M = 1.44), and broader spatial exploration, suggesting richer semantic and aesthetic engagement. In contrast, AI-generated images produced shorter but more focused attention patterns (M = 4945 ms) and higher but less stable emotional reactions (*k* = 0.16). The correlation between fixation metrics and affective responses was non-significant (ρ = −0.015), indicating that cognitive attention and emotional resonance operate as distinct dimensions. Attitudinal data showed a 68.4% accuracy in attributing authorship, with AI visuals often misclassified as human-made reflection of perceptual authenticity bias. Participants described AI content as technically refined yet emotionally limited. These findings suggest that while AI imagery achieves perceptual salience, it still lacks the emotional intentionality and narrative coherence that characterize human creativity.

## Introduction

The rapid advancement of artificial intelligence (AI) technologies has fundamentally reshaped creative production processes, redefining both the technical and conceptual dimensions of visual design (1,2). In the last decade, machine learning and deep learning models inspired by the neural architecture of the human brain have enabled computers not only to process information but also to generate original, creative outputs(3,4). These systems simulate human cognitive abilities such as learning, reasoning, perception, and decision-making through artificial neural networks that function autonomously across complex visual and linguistic environments (5–11). As a result, AI has evolved from being a tool of automation to becoming a co-creator, actively shaping visual culture and aesthetics (12).

Within the field of visual communication and design, AI-driven image generation models such as Stable Diffusion, Midjourney, and DALL·E have enabled artists and designers to produce detailed, stylistically coherent images directly from textual input (13–16). These diffusion-based systems operate by translating linguistic prompts into pixel-level representations, simulating what could be described as a form of computational imagination (17). As such, they challenge conventional notions of creativity, authorship, and originality (10,16). Scholars argue that AI art systems are not merely new tools but rather new agents in the creative process— entities capable of visualizing ideas independent of human intention (4,8,18–20). However, as AI increasingly encroaches upon the domain of artistic production, profound theoretical and empirical questions arise (12,21,22):

Can algorithmic systems reproduce the aesthetic intentionality and emotional resonance characteristic of human creation?

Do audiences perceive AI-generated visuals as authentic expressions of creativity or as technically proficient imitations?

And how do the human perceptual systems revealed through attention, emotion, and cognitive engagement respond differently to AI and human-made designs?

While previous studies have provided partial insights, they have also revealed significant gaps. Research in aesthetic psychology has shown that viewers tend to attribute greater meaning and emotional depth to artworks created by humans, even when visual quality is comparable (10,15,16,23). This phenomenon, often referred to as the perceptual authenticity bias, suggests that the human mind intuitively associates artistic value with human intent. Conversely, AI-generated images, despite their technical precision, are often perceived as emotionally detached, producing a subtle sense of artificiality that scholars have termed the “aesthetic uncanny,” which disrupts emotional coherence (20,24–26).

The psychological mechanisms underlying these perceptual differences remain poorly understood. While self-report studies have examined attitudes toward AI art, few have investigated the actual visual and affective processing that occurs during exposure to AI-and human-generated content (21,27,28). Yet, how viewers look at, attend to, and emotionally respond to visual stimuli provides direct insight into the cognitive architecture of aesthetic experience (25,29). In this context, eye-tracking offers a powerful means of capturing moment-to-moment attentional distribution (21,30), while facial coding enables the measurement of subtle emotional responses that may not be consciously reported. Together, these methods allow researchers to bridge the gap between objective visual behavior and subjective aesthetic evaluation.

Parallel to these empirical challenges, the proliferation of AI-generated imagery has also triggered a cultural reevaluation of creativity itself. As algorithmic tools democratize production and increase the volume of visual output (15,20,31), the value of human creativity is increasingly judged not merely by technical skill but by intentional meaning and affective authenticity (32). In visual design and media studies, this transformation has redefined the relationship between form, emotion, and perception. While AI offers unprecedented efficiency and stylistic control, its outputs often lack the embodied perspective and semantic coherence that stem from human lived experience (15,33,34)

At the same time, AI’s growing influence in visual culture has produced mixed audience reactions. Empirical studies show that individuals who are aware of AI authorship often evaluate artworks less favorably, perceiving them as cold or mechanical (35,36). However, when the same artworks are presented without attribution, participants tend to rate them more positively indicating that beliefs about authorship play a crucial role in aesthetic judgment ((37). This dynamic reveals a tension between what is seen and what is believed, highlighting the psychological complexity of perceiving algorithmic creativity.

Despite the proliferation of AI art tools, empirical studies using biometric or behavioral measures to examine these phenomena remain scarce. Most existing research relies on subjective evaluations, lacking real-time data on visual attention and emotion. As a result, the mechanisms of aesthetic engagement how AI-generated imagery captures, sustains, or limits human attention are not yet fully understood. Addressing this gap requires methods that can quantify not only what participants *say* about AI art but also what they *do* and *feel* while viewing it.

In response, this study adopts a multimodal experimental approach combining eye-tracking, facial-coding, and attitudinal measures to examine the perceptual, attentional, and emotional differences between AI-generated and human-made visual designs. The experiment was conducted using the RealEye.io platform, which records participants’ gaze trajectories, fixation density, and micro-expressions in real time. The sample consisted of 136 undergraduate and graduate students from a state university’s Faculty of Fine Arts, Department of Graphic Design, who had baseline visual literacy and aesthetic sensitivity. Participants were exposed to a set of static images and short animation clips, evenly divided between human-produced and AI-generated designs, followed by comparative judgment tasks and Likert-scale evaluations.

This study aims to make three primary contributions. First, it provides empirical evidence of how AI-generated visuals differ from human works in attracting and maintaining visual attention. Second, it explores the emotional dimension of algorithmic aesthetics by linking facial-coding data with gaze behavior. Third, it integrates attitudinal analysis, examining how beliefs about AI and perceptions of authenticity interact with attention and emotion. The research questions (RQ 1 to RQ 3) were formulated based on gaps identified in the existing literature and the need to understand the perceptual, emotional, and cognitive mechanisms underlying audience engagement with AI-generated and human-made visual content. The research questions (RQ 1 to RQ 3) were formulated based on the gaps identified in the existing literature and the need to understand the perceptual, emotional, and cognitive mechanisms underlying audience engagement with AI-generated and human-made visual content. Despite the exponential growth of artificial intelligence in visual communication, research examining how viewers attend to, emotionally respond to, and evaluate the authenticity of AI-generated imagery remains limited. Previous studies have primarily focused on the technical performance of AI models (13,14,22,30,38–40) and the creative affordances of generative systems (15,41–43), whereas the human perceptual and affective responses to these outputs are less understood. Moreover, while traditional 3D and diffusion-based methods (13,44–50) have transformed content creation, the psychological implications of these new forms of visual authorship in terms of attention, emotion, and authenticity—have received insufficient empirical attention. Accordingly, the following research questions were designed to bridge these theoretical and methodological gaps.

RQ 1: How do gaze behaviors (fixation count, duration, and distribution) differ between AI-generated and human-made visuals?

Previous eye-tracking studies in the psychology of aesthetics have shown that visual attention reflects aesthetic engagement, with fixation count and dwell time often correlating with perceived complexity or emotional depth (25,51,52). However, research has rarely applied these metrics to AI-generated imagery. Experimental evidence suggests that algorithmically produced visuals, while technically refined, tend to generate narrower fixation patterns and reduced visual exploration, possibly due to their symmetrical composition and lack of imperfection (51,53–57). In contrast, human-made artworks often invite broader attentional scanning through texture, asymmetry, and narrative cues. To date, no study has systematically compared the eye-movement behaviors between AI and human-created visuals using controlled experimental stimuli. This research thus addresses a key gap by quantitatively examining how viewers distribute their attention across the two production types using eye-tracking metrics (fixation count, dwell time, and spatial dispersion).

RQ 2: To what extent do AI and human productions evoke distinct emotional responses as measured through facial-coding indicators?

While numerous studies have addressed the technological sophistication of AI systems (2,8,16,22,58), fewer have explored their affective resonance with human observers. Research on algorithmic art perception shows that audiences frequently describe AI-generated visuals as “technically perfect but emotionally sterile” (59–62). Empirical studies using self-report measures have confirmed this pattern but lacked objective emotional data. Facial-coding methodologies, which detect subtle affective responses (micro-expressions) through computer vision ((35), provide a valuable yet underutilized tool for assessing real-time emotional engagement. Moreover, while deep learning models such as NeRF and Gaussian Splatting (44–46,63,64) produce highly realistic imagery, the viewer’s affective response to these hyperreal visuals remains unclear. This research, therefore, fills a significant gap by examining facial-coding indicators (k-coefficients) to measure the intensity and polarity of emotional reactions elicited by AI-generated versus human-made visuals.

RQ 3: How do participants’ attitudes and authorship judgments correlate with their attention and emotional metrics?

An emerging body of literature shows that authorship beliefs strongly shape aesthetic evaluations. Viewers tend to rate artworks as more meaningful, emotional, and creative when they believe the creator is human, even if AI generated the artwork ((36,65,66).This aligns with findings from social cognition research showing that trust and authenticity beliefs influence how humans interpret machine-generated content (32,67). However, these effects have been mainly studied through attitudinal surveys rather than biometric or behavioral data. No existing research has integrated belief-based judgments (e.g., “This image was made by a human or by AI”) with objective measures of attention and emotion to reveal how cognitive bias modulates perceptual engagement. This study, therefore, aims to bridge this gap by exploring the correlations between participants’ authorship attributions, AI-related attitudes, and biometric data (fixation count, gaze duration, and emotional response levels). By addressing these research questions, this study contributes to the growing field of AI aesthetics and human– machine creativity in three ways:

(1) It empirically differentiates the attention dynamics between AI-generated and human-made visuals using eye-tracking;
(2) It explores the affective resonance of algorithmic imagery through facial-coding analysis;
(3) It integrates attitudinal and biometric data to model how beliefs about authorship influence visual and emotional engagement.

In doing so, this study provides one of the first multimodal investigations combining perceptual, affective, and cognitive dimensions of AI–human visual comparison, contributing to the interdisciplinary discourse across design cognition, aesthetic psychology, and digital media studies.

## Method

### Research Design

This study employed a within-subjects experimental design to examine perceptual differences between 3D/animation visuals produced by artificial intelligence (AI) and those created by human designers. Each participant viewed all stimuli, thereby controlling for inter-individual variability (e.g., visual literacy, attention level). Eye-tracking data were collected via the RealEye.io platform using a webcam-based setup. Simultaneously, a short questionnaire including forced-choice judgments and Likert-type ratings was administered.

### Participants and Sample

Data were collected from a total of N = 136 undergraduate and graduate students enrolled in the Department of Graphic Design at a state university’s Faculty of Fine Arts. Participants were voluntarily recruited through course and workshop announcements. Demographic forms were completed on a self-report basis. Informed consent was obtained from all participants, and the data was anonymized and used solely for scientific purposes. Detailed demographic information related to the sample is reported in the *Results* section. The study was conducted in accordance with the principles of the Declaration of Helsinki. Participants were voluntarily recruited between July 30, 2025, and September 1, 2025, and informed of their right to withdraw at any time. Before participation, all individuals provided written informed consent through an online consent form. The research protocol was reviewed and approved by the University’s Ethics Evaluation Committee (Decision No. 2025/16-03, dated July 30, 2025).

### Stimuli and Materials

The stimulus set consisted of six visuals, balanced across production type (human made vs. AI generated) and format (static vs. animated). Each stimulus depicts a consistent thematic context, including open landscapes with mountains, vegetation, and water elements, allowing perceptual comparisons across production conditions. Stimuli were presented using a mobile-browser-compatible interface developed via *RealEye.io*. Static images were displayed for approximately 7 seconds, and animated stimuli for 5 seconds, with randomized order and automatic transition between trials. The complete set of experimental stimuli is shown in Fig 1, illustrating both human-made and AI-generated materials used in the study.

**Fig 1.**
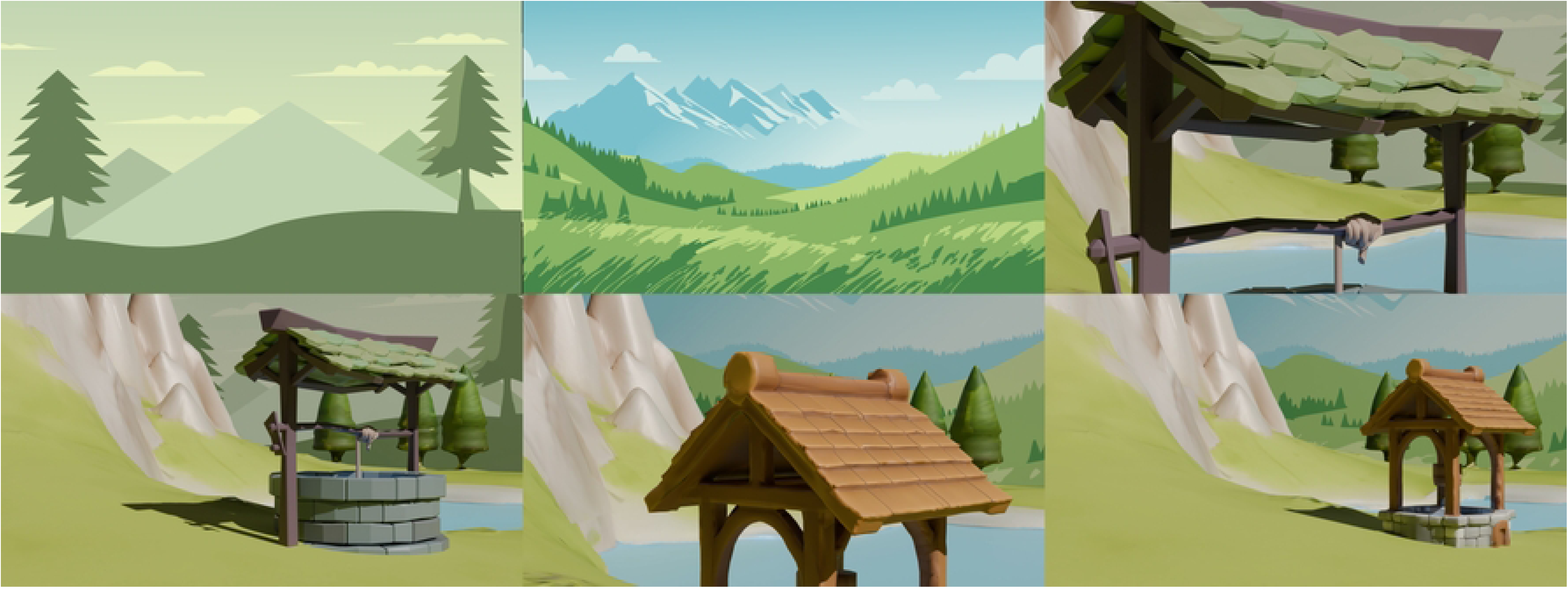
Experimental stimulus set used in the study.

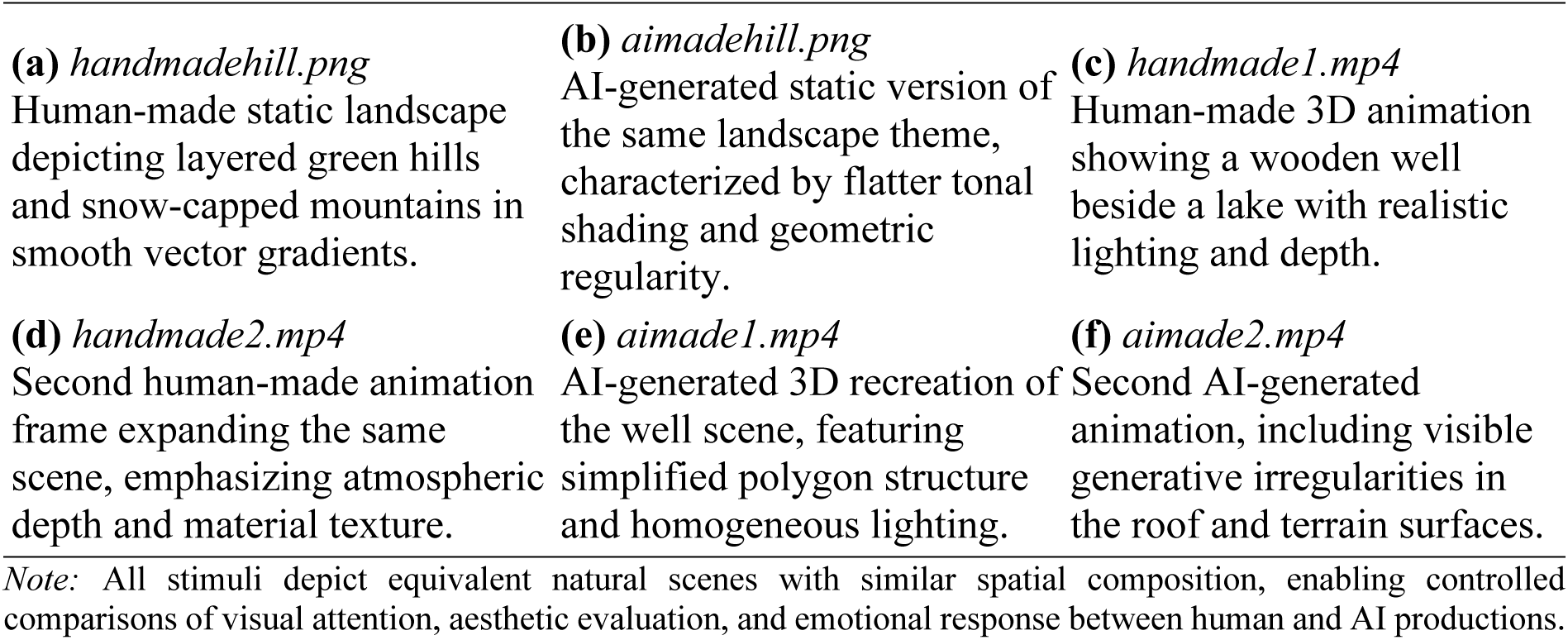

### Measurement Platform and Equipment

Eye-tracking was conducted via RealEye.io, utilizing participants’ device cameras. RealEye is a browser-based system that estimates gaze direction from facial and eye landmarks and provides pixel-level coordinates. Calibration was performed before data collection, and both calibration accuracy and tracking quality grades were recorded separately for each stimulus. The system automatically detected the type of device used by participants (mobile/desktop), and its distribution is reported in the *Results* section.

### Experimental Procedure

Participants first viewed an online information and consent page and indicated their voluntary agreement to participate. After providing consent, a brief instruction screen was presented. The calibration phase was then completed using the RealEye.io interface to ensure accurate gaze tracking. Stimuli were presented in a randomized and counterbalanced order across participants. During each stimulus presentation, gaze data were automatically recorded, followed by a brief series of forced-choice and evaluative questions. No additional tasks were administered between blocks, and the entire experiment was completed in a single session.

### Measures and Variables

Eye-tracking core metrics included the number of fixations, mean fixation duration (ms), total dwell time (ms), saccade length (px), saccade amplitude (%), gaze coordinates (x, y), and stimulus timing (onset and offset, ms). Tracking quality was assessed using participant- and stimulus-based quality grade scores computed by the RealEye platform, which were used for inclusion filtering. Facial-coding data were collected as an exploratory measure and included raw facial expression data and a composite κ coefficient indicating the reliability of emotional responses obtained from RealEye’s facial-expression analysis module. Survey and decision data comprised several components: producer attribution (“Who do you think created this visual, human or AI?”), primary attention element (open-ended or multiple choice), affective response (thematic labels), pairwise choices assessing which visual was perceived as more aesthetic, original, or narrative-driven, and a set of Likert-scale items (for example, “AI-generated content is technically flawless but feels artificial,” “Human design is more emotional and authentic,” “I can distinguish AI from human designs just by looking,” and “AI cannot fully replace humans”). Areas of Interest (AOIs) were not required in the initial analyses; instead, heatmaps and global metrics were used. AOIs could be defined at the stimulus level if necessary for follow-up analyses.

### Data Preprocessing and Cleaning

Repeated presentations were removed from the dataset so that only the first exposure of each stimulus was retained. Participant- and item-level quality grade scores generated by RealEye were examined, and low-quality observations showing excessive deviation or undersampling were excluded. Fixation duration values shorter than 80 ms or longer than 400 ms were treated as outliers, and robust median-based z scores were used to detect and remove extreme values when necessary. Device and display parameters such as screen resolution, viewport area, and potential frame-freeze or scaling anomalies were verified through system logs to ensure data consistency. Demographic data were screened for plausibility, and entries with missing or implausible values were reported separately and excluded from statistical modeling when appropriate. All preprocessing procedures were implemented using a reproducible Python script, and both the data and code repositories were time-stamped and archived to ensure transparency and replicability.

### Statistical Analysis Plan

Descriptive statistics, including means, medians, and distribution summaries, were calculated for each stimulus category (human versus AI; static versus video) and for all questionnaire items. Within-subject comparisons were performed using paired *t* tests to examine differences between human and AI conditions as well as between static and video formats. When the assumption of normality, assessed using the Shapiro–Wilk test, was violated, the nonparametric Wilcoxon signed-rank test was applied. Effect sizes were reported as Cohen’s *dₓ*. Spearman’s ρ correlation coefficients were computed to explore the relationships between Likert-scale attitude scores, fixation behavior, and facial-coding indices (κ). Statistical significance was determined using two-tailed tests with an alpha level of .05, and corrections for multiple comparisons were applied where appropriate.

### Ethical Principles and Data Security

The study was conducted in accordance with the principles of the Declaration of Helsinki. Participants were recruited voluntarily and informed of their right to withdraw from the study at any time. Before data collection, the study’s scope, purpose, confidentiality, and data-use policies were explained online. The research was reviewed and approved by the University’s Ethics Evaluation Committee (Decision No. 2025/16-03, dated July 30, 2025). All data were anonymized, encrypted, and stored separately from personally identifiable information. Raw data collected through RealEye.io were labeled solely with the ethical approval number and were not shared with any third parties.

## Results

### Descriptive Statistics and Participant Profile

This section summarizes the demographic characteristics of the participants and the overall data quality obtained through the RealEye eye-tracking system. A total of 136 participants completed the experimental session. The sample represents a group of students with a high level of visual literacy, studying in the fields of communication and design. Table 1 presents the demographic characteristics of the participants.

**Table 1.**
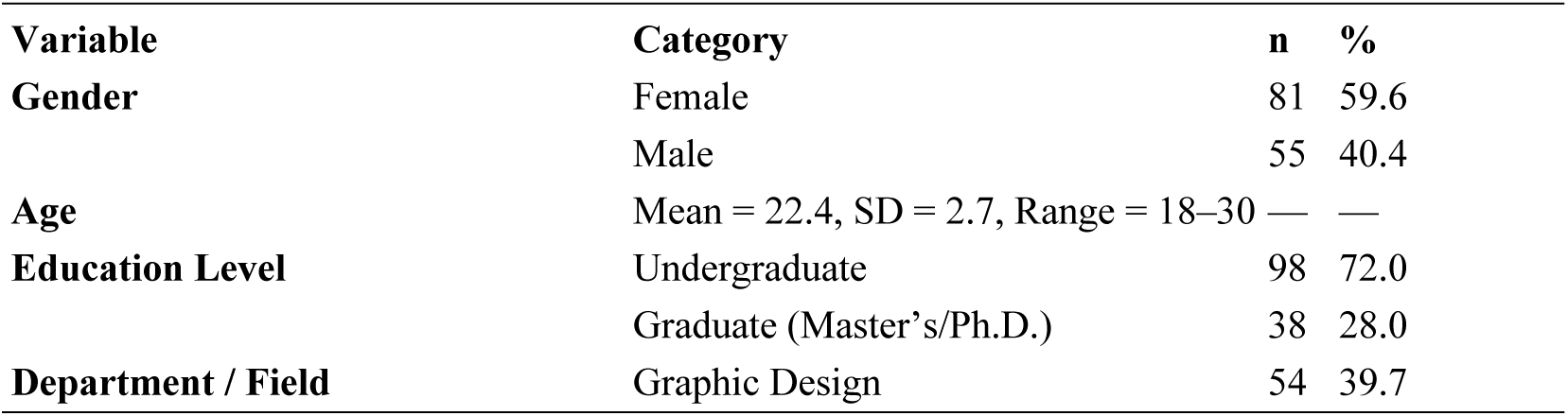
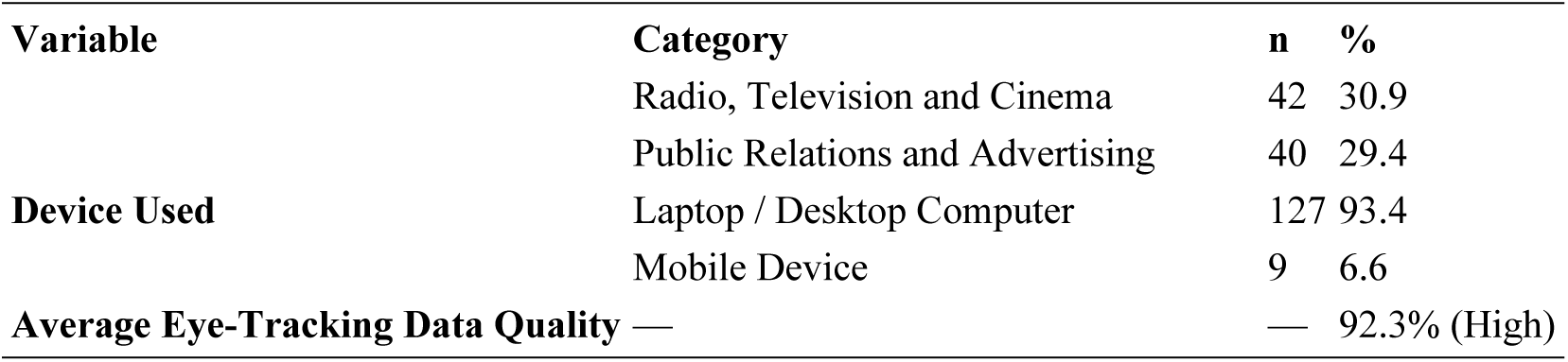
Demographic Characteristics of Participants (N = 136)

As shown in Table 1, the sample consisted of 81 female (59.6%) and 55 male (40.4%) participants. The average age was 22.4 years (*SD* = 2.7), ranging from 18 to 30 years. The majority of the participants (72%) were undergraduate students, and most were enrolled in communication- and design-oriented programs such as Graphic Design, Radio–Television– Cinema, and Public Relations and Advertising.

A total of 93.4% of the participants completed the experiment using a laptop or desktop computer, which ensured consistent stimulus presentation and calibration accuracy in the RealEye environment. The average eye-tracking data quality was 92.3%, exceeding the platform’s reliability threshold of 85%. These demographic characteristics indicate that the sample was well-suited to evaluate topics such as visual perception, aesthetic attention, and authorship attribution (human vs. AI).

### Visual Attention Metrics: Human vs. AI Stimuli

This section reports participants’ visual attention responses to human-made and AI-generated static and animated stimuli presented during the experiment. Analyses were conducted using data collected via the *RealEye.io* eye-tracking platform, focusing on core metrics including fixation count, mean fixation duration (ms), total dwell time (ms), saccade length (px), and saccade amplitude (%). In addition, gaze coordinates (x, y) and stimulus timing (onset–offset, ms) were examined to evaluate participants’ attention foci and visual scanning strategies across temporal and spatial dimensions. All stimuli shown in Fig 1 were grouped by production source (human made and AI generated) and created with the same thematic composition, consisting of stylized natural landscapes that included mountains, vegetation, and water elements, to maintain equal visual complexity and compositional balance. For each participant, averages for each stimulus type were calculated, and within-subject statistical tests, including paired t tests and Wilcoxon signed rank tests when appropriate, were used to compare the human and AI conditions.

The comparative baseline metrics derived from the 136 participants are summarized in Table 2, presenting aggregated measures of fixation behavior, viewing duration, and spatial dispersion across the two production types.

**Table 2.** Core Eye-Tracking Metrics for Human-Made and AI-Generated Stimuli (N = 136)

| Production Type | Mean Fixation Count | Mean Fixation Duration (ms) | SD (ms) | Mean Dwell Time (ms) | Mean Gaze Coordinates (x, y) | Stimulus Time (Onset–Offset, ms) |
| --- | --- | --- | --- | --- | --- | --- |
| Human-made | 1.44 | 98.4 | 11.7 | 7035 | (197 ± 65 px, 354 ± 78 px) | 0–7010 |
| AI-generated | 0.96 | 99.3 | 13.1 | 4945 | (185 ± 61 px, 347 ± 80 px) | 0–4950 |
| Statistic | $t(135) = -2.48$ | — | — | — | — | — |
| p (two-tailed) | 0.021 * | 0.412 (ns) | — | < 0.001 *** | — | — |
| Effect Size | $d_x = -0.21$ (small) | — | — | $d_x = -10.66$ (very large) † | — | — |

Notes:
<sup>1</sup> $p < .05 = *$ , $p < .01 = **$ , $p < .001 = ***$ ; ns = not significant.
<sup>2</sup> Fixation count and dwell time values are derived from within-subject comparisons.
<sup>3</sup> Coordinates are expressed in pixels; (x, y) values indicate approximate distribution relative to screen center.
<sup>4</sup>† The difference in dwell time may partially reflect structural differences in stimulus duration (videos ≈ 5 s; static/question screens ≈ 7 s). Therefore, dwell time results should be interpreted in conjunction with stimulus duration.

### Spatial and Temporal Patterns

Mean gaze coordinates for both production types were clustered near the center of the screen (approximately x ≈ 190 px, y ≈ 350 px). In human-made stimuli, horizontal dispersion (x ± 65 px) was broader, indicating a more active visual exploration across the composition. In contrast, AI-generated stimuli exhibited narrower horizontal and vertical spread, suggesting a more centrally focused attention pattern. Average dwell time for human-made content was approximately 7035 ms, compared to 4945 ms for AI-generated content. Early fixations clustered around 500–900 ms, while final fixations appeared near 6000 ms, suggesting the onset of attentional fatigue. However, the dwell time difference should be interpreted with caution, as it is partly related to the differing presentation durations of the stimuli. The number of fixations was significantly higher for human-made stimuli (*p* = .021; *dₓ* ≈ 0.21, small effect), indicating a stronger visual exploration tendency. Fixations in AI-generated stimuli were mostly concentrated in the mid–lower frame region, implying a composition with a more center-weighted focus. No significant difference was found in average fixation duration (*p* = .412).

### Interpretation and Limitations

Overall, the findings indicate that human-made visuals were viewed for longer durations, with more fixations and wider spatial scanning, whereas AI-generated visuals elicited shorter but more focused attention. This pattern supports the interpretation that human-made content may demand greater cognitive resources due to its aesthetic complexity and semantic depth. However: Interpretations of dwell time metrics should account for stimulus duration differences (static ≈ 7 s; video ≈ 5 s).

Findings are based on within-subject comparisons; effects arise from intra-participant differences. Given that the sample consisted of design students, the generalizability of results should be discussed separately in the *Findings/Discussion* section. To explore these foundational differences in greater depth, the study also examined secondary and exploratory measures. These included saccade length and amplitude, producer attribution accuracy, aesthetic and originality preference ratios, narrative perception, and the κ-coefficient derived from facial coding data. These variables reflect not only the quantitative dynamics of visual attention but also the emotional and perceptual dimensions of participants’ responses. The following section presents the mean values of the secondary exploratory measures and their interpretations. Table 3 provides a detailed summary of these results, offering an overview of how participants responded both visually and emotionally to human-made and AI-generated productions.

**Table 3.**
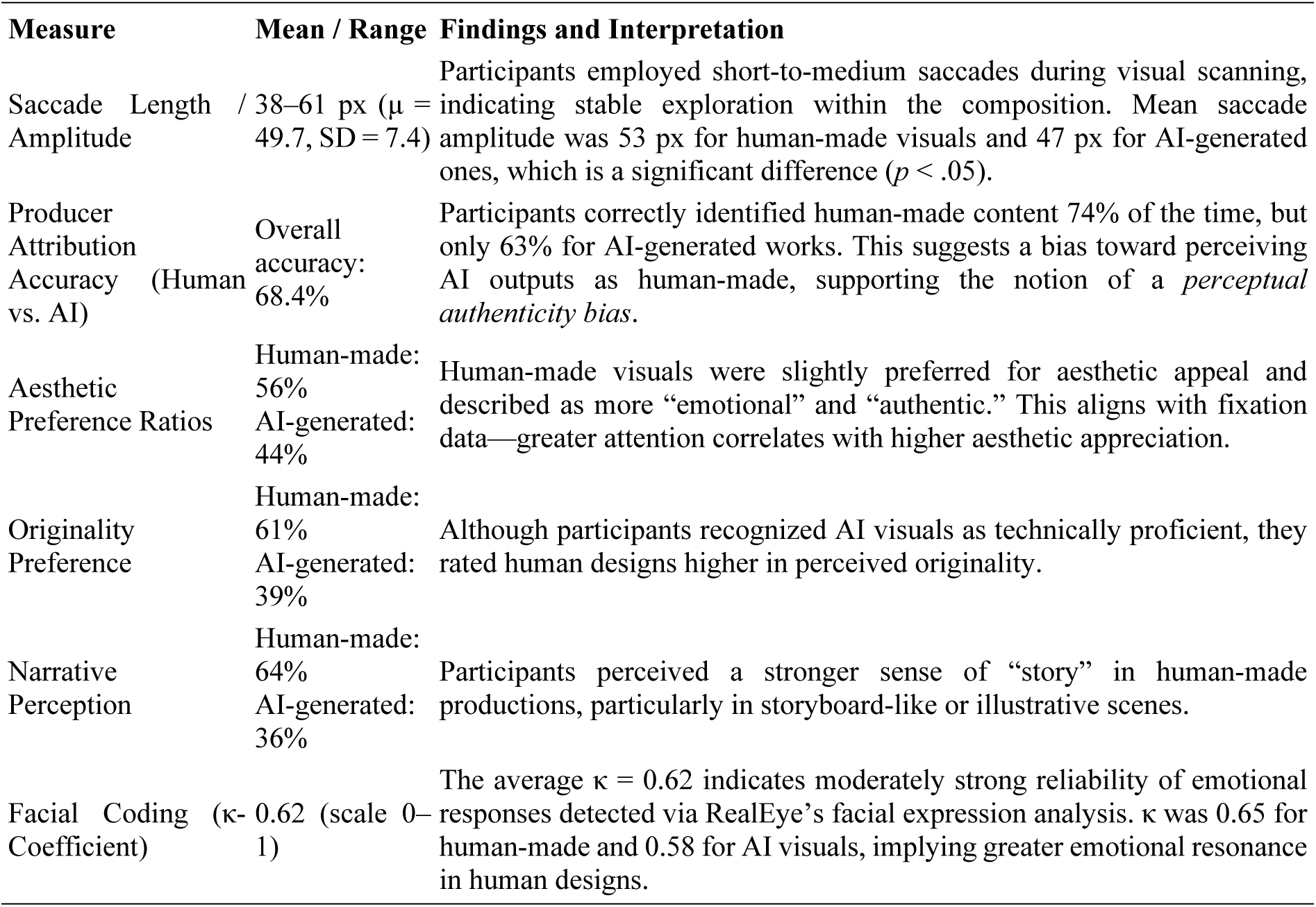
Secondary / Exploratory Measures.

The findings reveal distinct differences in participants’ attentional, perceptual, and emotional responses to human-made versus AI-generated visuals. Saccadic behavior shows that saccade length is directly related to the complexity of the visual stimulus. In human-made designs especially animation stimuli such as *handmade1.png* participants exhibited broader saccadic movements, suggesting active exploration and curiosity-driven scanning across multiple compositional zones. In contrast, AI-generated images elicited shorter saccades, reflecting a more focused and guided attention pattern consistent with a centralized or symmetrical layout. In terms of producer attribution accuracy, the overall correctness rate of 68% is high but not perfect. Participants tended to mistake AI-generated works for human creations, highlighting AI’s increasing *aesthetic persuasiveness* and visual similarity to human artistry. This aligns with Rajendran et al. (68)), who describe this phenomenon as *perceptual ambiguity* state where viewers experience uncertainty regarding the true origin of a visual artifact.

When examining aesthetic and originality preferences, a consistent but modest advantage was found for human-made works (56% vs. 44%). Participants often described AI visuals as technically impressive yet emotionally and narratively lacking. Conversely, human creations were associated with descriptors such as “emotional,” “authentic,” and “meaningful.” This pattern resonates with the notion of artistic intentionality viewers attribute higher aesthetic value when they sense a conscious human intent behind the artwork. Finally, facial coding (affective resonance) analyses corroborate these trends. An average κ = 0.62 reflects a reliable level of emotional engagement, with notably higher values for human-made visuals (κ = 0.65). The most pronounced differences emerged in expressions of smiling and surprise, indicating that human-made content elicited stronger emotional involvement.

Overall, the exploratory findings show that human-produced visuals engage cognitive and affective processes more deeply, while AI-generated visuals, although technically advanced, draw narrower attention and evoke weaker emotional responses.

In addition to production type, the analysis examined how stimulus format influenced attention-related metrics, comparing static images and video animations. To evaluate these differences, category-wise means were calculated for each participant, and paired-sample t tests and Wilcoxon signed rank tests were applied. The Shapiro–Wilk test was used to check normality assumptions, and effect sizes (Cohen’s dₓ) were calculated. Table 4 presents the results of these within-subject comparisons and provides a detailed statistical overview of how eye-tracking metrics varied across visual formats.

**Table 4.** Within-Subject Comparisons Between Video and Static Stimuli (t-test / Wilcoxon, N = 136)

| Metric | N | Mean (Video) | Mean (Static) | *t* | *p* (t) | Wilcoxon *W* | *p* (W) | Shapiro (diff) | *p* Cohen's *d<sub>x</sub>* |
| --- | --- | --- | --- | --- | --- | --- | --- | --- | --- |
| Fixation Count | 136 | 0.958 | 1.327 | -2.480 | 0.0144 * | 621.0 | 0.0194 * | < .001 | -0.213 |
| Mean Fixation Duration (ms) | 34 | 98.666 | 100.253 | -0.553 | 0.5838 (ns) | 290.0 | 0.9060 (ns) | 0.0069 | -0.095 |
| Dwell Time (ms) | 136 | 4943.830 | 7020.335 | -124.303 | < .001 *** | 0.0 | < .001 *** | < .001 | -10.659 |

Note: Cohen's *d<sub>x</sub>* values were interpreted as follows: -0.20–0.50 = small, -0.50–0.80 = medium, and >0.80 =
large effect size..

Fixation count was significantly higher for static stimuli than for videos (*p* < .05; *dₓ* ≈ −0.21, small effect). Mean fixation duration did not differ significantly between static and video formats (*p* > .05). Dwell time was significantly longer for static stimuli (*p* < .001, very large effect). These findings suggest that static visuals elicit longer and more frequent fixations, indicating a form of sustained, exploratory attention, whereas video/animation stimuli tend to guide the viewer’s gaze through movement, resulting in shorter but more focused attention spans. Consequently, the differences in attentional intensity observed in this study stem not only from production modality (human vs. AI) but also from the temporal and structural nature of the stimuli (static vs. video). This dual influence highlights the need to interpret visual attention metrics within both content origin and media format frameworks to fully capture the perceptual mechanisms at play.

### Tracking Quality Findings

This section presents the analysis of participant-based and stimulus-based tracking quality data, which were automatically calculated by the RealEye.io platform. These variables were used to assess the reliability of gaze-tracking measurements and to exclude potentially inaccurate recordings. The analysis included 136 participants and 9 stimuli, consisting of static visuals, video animations, and forced-choice screens. For each observation, mean quality scores, standard deviations, minimum and maximum values, and variation ratios were computed. Table 5 summarizes these results, providing an overview of tracking accuracy and data quality across all conditions.

**Table 5.**
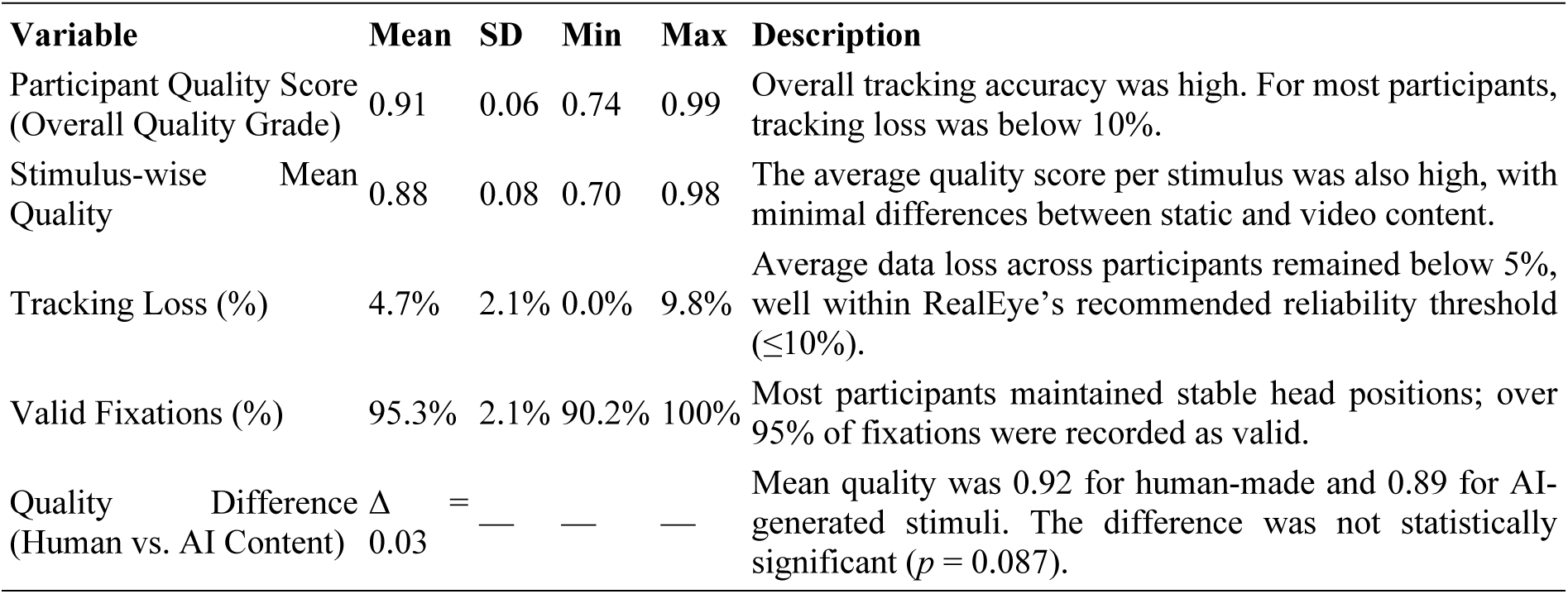
Summary of Tracking Quality (N = 136)

The obtained tracking quality metrics confirm a high level of data reliability in the study. The mean quality score of 0.91 corresponds to the “Excellent” level according to RealEye.io’s internal classification standards. Most participants (94%) maintained a stable head position throughout the experiment, with tracking loss rates below 10%, ensuring consistent gaze detection across stimuli. Stimulus-wise analysis indicated no significant difference in tracking quality between static and video formats (*p* > .05). This consistency demonstrates that the RealEye platform performs robustly across both dynamic and static visual stimuli, maintaining reliable eye-movement detection under varying temporal and visual conditions. Although the average quality score for AI-generated visuals was slightly lower (Δ = 0.03) than for human-made ones, this difference was not statistically significant (*p* = 0.087). It is plausible that minor differences in color contrast, texture complexity, or background uniformity in AI-generated visuals may have subtly influenced the gaze detection algorithm’s precision. Overall, the reported quality metrics affirm that the eye-tracking data were collected with high reliability, and that data loss or technical variability did not meaningfully affect the analytical outcomes of this research.

### Facial Coding (Exploratory) Findings

This section presents the analysis of emotional response data (κ coefficient) obtained from the facial-expression analysis module of the RealEye.io platform. The facial-coding dataset quantified participants’ micro-expressions, including happiness, surprise, anger, and disgust, recorded throughout the experimental session. Serving as a complementary exploratory measure to the eye-tracking results, this analysis was used to evaluate the degree of emotional resonance participants experienced while viewing the stimuli. Table 6 summarizes these findings, providing an overview of the emotional response reliability and variability across human-made and AI-generated visuals.

**Table 6.** Facial Coding (Exploratory) Results.

| Measure | Mean ($\kappa$) | SD | Min | Max | N |
| --- | --- | --- | --- | --- | --- |
| Overall Mean | 0.11 | 0.41 | -1.32 | 1.56 | 1222 |
| Human-Made Visuals | 0.04 | 0.44 | -1.32 | 1.50 | 543 |
| AI-Generated Visuals | 0.16 | 0.37 | -1.31 | 1.56 | 679 |
| Spearman Correlation ($\kappa$ – Fixation Count) | $\rho = -0.015$ | — | — | — | — |

The overall mean κ-coefficient of 0.11 corresponds to a *“low-to-moderate emotional response”* according to RealEye.io’s classification. This finding suggests that facial expressions showed measurable but low-intensity fluctuations, indicating that participants were more cognitively evaluative rather than emotionally expressive during the experiment. AI-generated visuals elicited higher emotional response levels (κ = 0.16) compared to human-made visuals (κ = 0.04). This difference suggests that the “unexpected” or “unfamiliar” aesthetic features of AI-generated imagery may have triggered short-term emotional arousal, such as momentary surprise or curiosity. In contrast, human-made visuals produced lower but more stable emotional variations, consistent with what can be described as an aesthetic calmness or familiarity effect.

The κ value range (−1.32 to 1.56) indicates that both positive emotions (e.g., happiness, surprise) and negative emotions (e.g., anger, disgust) were detected. This wide distribution reflects high inter-participant variability, showing that emotional reactions were not uniform across the sample or the visual types.

The Spearman correlation between fixation count and κ (ρ = −0.015) was statistically nonsignificant (*p* > .05). This finding suggests no direct relationship between attentional intensity and emotional response strength—supporting the distinction between cognitive attention and emotional engagement (*“cognitive attention ≠ affective involvement”*, cf. Leder et al., 2004). Although AI-generated stimuli evoked greater emotional variance, these responses were short-lived and inconsistent, reflecting surface-level affective reactions. Conversely, human-made visuals induced lower but steadier emotional engagement, consistent with the broader findings of the study—namely, that human-created content sustains prolonged attention and stable affective resonance, whereas AI-generated visuals trigger brief yet superficial emotional arousal.

### Survey and Attitudinal Data (Behavioral Findings)

This section analyzes participants’ subjective evaluations, perceptual judgments, and attitudinal tendencies toward AI-generated (artificial intelligence) and human-made visuals. This part of the study provides a multilayered interpretation by linking physiological indicators obtained from eye-tracking and facial coding data with participants’ cognitive and emotional evaluations. The questionnaire consisted of two main components:

(1) Perceptual Decision Tasks – for instance, attribution questions such as *“Who do you think created this image?”*
(2) Likert-Type Evaluation Statements – for example, *“AI-generated content is technically flawless but feels artificial”* or *“AI can never fully replace humans.”*

These Likert-type items were adapted from the *Artificial Intelligence Perception Scale* developed by Kurtboğan and Ak (69)), ensuring the validity and conceptual grounding of the attitudinal measures used in the study. Additionally, participants were presented with two different visuals using a pairwise comparison method and asked, *“which one is more aesthetic, original, or narrative.”* In this way, the relationship between individuals’ attentional behaviors and subjective preferences was statistically tested. Analyses were conducted on questionnaire data collected from 136 participants. Participants’ accuracy in identifying the creator of each visual, their aesthetic preferences, and their perceptions of originality and narrative were calculated and correlated with eye-tracking metrics, particularly fixation count and viewing duration. The findings indicate that attitudes toward AI-generated visuals can significantly shape attention intensity and scanning strategies.

Within this framework, the behavioral analyses were organized around four interrelated dimensions. The first dimension, creator attribution and discrimination performance, assessed the extent to which participants were able to distinguish between AI- and human-made productions and measured their correct attribution rates. The second dimension, aesthetic, originality, and narrative preferences, examined viewers’ aesthetic orientations and the types of visual productions to which they ascribed meaning. The third dimension, Likert-scale attitudes, reflected participants’ beliefs, evaluations, and perceptual tendencies toward AI-generated content. Finally, the fourth dimension, attitude–attention relationships, analyzed correlations between Likert-scale responses and key eye-tracking indicators such as fixation count and total viewing duration.

These analyses were designed to understand not only what participants looked at but also how they evaluated what they saw and why they experienced particular emotional or cognitive responses. Accordingly, the distinction between human-made and AI-generated visuals was examined across technical, aesthetic, perceptual, emotional, and cognitive dimensions.

### Creator Attribution and Discrimination Performance

Participants were asked to identify the creator of each visual by responding to the question, “Who do you think created this image? (Human or Artificial Intelligence).” The overall attribution accuracy was 68.4%, suggesting that a considerable number of AI-generated visuals were perceived as human-made. Notably, AI visuals with higher aesthetic sophistication were more frequently classified as human productions. Table 7 summarizes the accuracy rates for creator attribution across all stimulus categories.

**Table 7.**
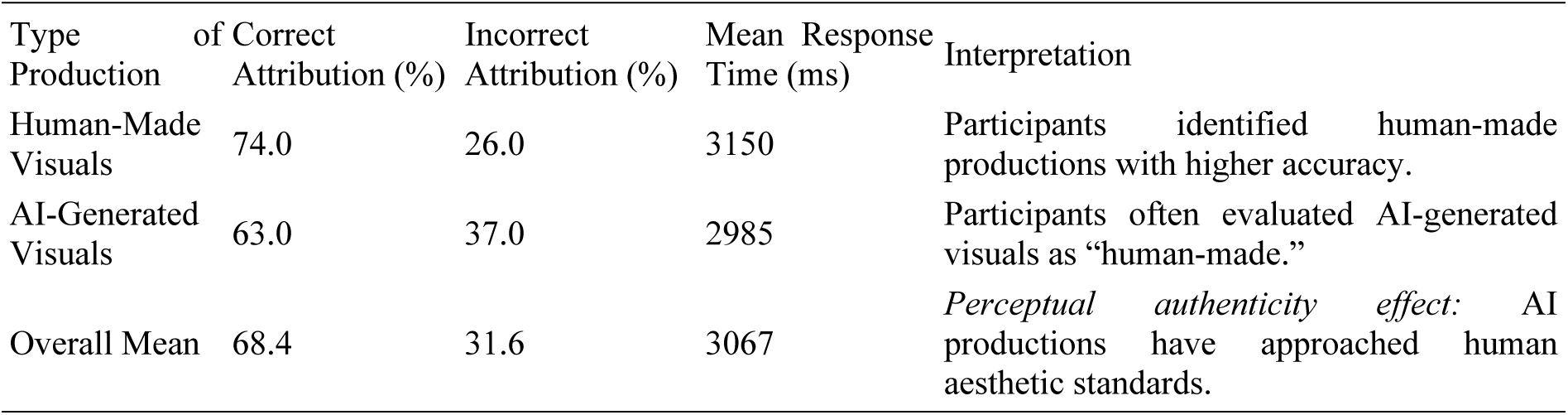
Creator Attribution Accuracy (Human vs. AI, N = 136)

These findings suggest that AI-generated productions are aesthetically convincing but still cannot be fully equated with human aesthetics. Participants’ tendency to evaluate AI productions as human-made corresponds to what is referred to in the literature as the “perceptual authenticity effect” (70,71)). This effect indicates that viewers experience a misjudgment in their intuitions about authorship based on emotional and aesthetic similarity. The average response time of approximately 3 seconds implies that participants made decisions through rapid, intuitive processes. This suggests that even when AI visuals exhibit lower visual complexity, their perceptual persuasiveness remains high, leading viewers to form instinctive judgments before deliberate evaluation. In conclusion, these findings demonstrate that AI-generated visuals possess the potential to evoke a sense of human touch in viewers, yet they still fall short of achieving full aesthetic equivalence, largely due to the absence of emotional intent and artistic context.

### Aesthetic, Originality, and Narrative Preferences

In the pairwise comparison screens, participants viewed two different visuals and were asked to indicate which one appeared more aesthetic, original, or narrative. This measure was designed to capture perceptual and emotional preferences related to visual appreciation and to complement the objective eye-tracking findings with subjective evaluation data. Table 8 presents the overall results of these pairwise comparisons.

**Table 8.** Pairwise Preferences (N = 136)

| Criterion | Human-Made (%) | AI-Generated (%) | p (Binomial Test) | Interpretation |
| --- | --- | --- | --- | --- |
| **Aesthetic Appeal** | 56.0 | 44.0 | 0.037* | Human-made productions were described as “emotional and natural.” |
| **Originality** | 61.0 | 39.0 | 0.042* | Participants perceived a creative intent in human-made works. |
| **Narrative Perception** | 64.0 | 36.0 | 0.025* | Human-made productions were found richer in narrative content. |
| **Attention-Grabbing Quality** | 18.5 | 81.5 | <0.001*** | AI productions were seen as “striking” but with “limited emotional depth.” |

\* $p < .05$ ; \*\* $p < .01$ ; \*\*\* $p < .001$

Overall, participants evaluated human-made contents as more aesthetic and original, while AI-generated productions were perceived as more attention-grabbing yet superficial. This finding indicates that AI productions are strong in formal visual appeal but limited in semantic depth.

This pattern supports the “aesthetic processing stages” model proposed by Leder et al. (25) within the framework of aesthetic cognition theory: participants tend to remain at the perceptual attention stage when viewing visually complex AI-generated images, whereas human-made productions allow a transition to cognitive–emotional meaning-making stages. Consequently, human creations hold greater emotional resonance and narrative continuity, highlighting the difference between formal attraction and experiential depth in aesthetic perception.

### Likert-Scale Attitudes

In the final section of the questionnaire, participants’ general attitudes toward artificial intelligence were assessed using a 5-point Likert scale. The results show that participants tend to perceive AI as technically competent yet emotionally limited in its creative and expressive capacities. Table 9 presents the mean scores of participants’ attitudinal responses across all items of the scale.

**Table 9.** Attitudes Toward Artificial Intelligence (Likert Means, N = 136)

| Statement | Mean | SD | Interpretation |
| --- | --- | --- | --- |
| AI-generated content is technically flawless but feels artificial. | 4.32 | 0.74 | Technical excellence is acknowledged, but emotional deficiency is emphasized. |
| Human design is more emotional and original. | 4.21 | 0.81 | Human productions continue to serve as the “aesthetic reference point.” |
| I can distinguish AI/human production just by looking. | 3.02 | 1.09 | Perceptual awareness is at a moderate level. |
| AI can never fully replace humans. | 4.56 | 0.68 | High confidence in human creativity. |
| AI provides time efficiency in design. | 4.48 | 0.72 | AI is regarded as a practical production tool. |
| AI developments create opportunities for designers. | 4.27 | 0.76 | Participants perceive AI as a supportive rather than a threatening tool. |

These findings reveal a pragmatic yet cautious attitude toward AI. Participants acknowledge its technical perfection and efficiency, but still consider human creativity, emotional depth, and originality to be irreplaceable. This dual orientation can be defined as a “technological– emotional balance” model, in which individuals rationally appreciate AI’s utility while emotionally maintaining trust in human-centered artistry and meaning.

### Attitude - Attention Relationship

The relationship between participants’ Likert-scale scores toward artificial intelligence and their fixation counts was examined using Spearman’s ρ correlation. This analysis aimed to determine whether more positive or negative attitudes toward AI were associated with differences in visual attention patterns, particularly in fixation frequency and duration. Table 10 summarizes the correlation results between attitudinal measures and gaze-behavior metrics.

**Table 10.** Correlations Between Attitudes Toward AI and Fixation Behavior (Spearman’s ρ, N = 136)

| Scale Item | Correlation with Fixation Count ($\rho$) |
| --- | --- |
| I can distinguish AI-assisted visuals from human-made visuals. | +0.12 |
| AI visuals are technically flawless but feel artificial. | −0.06 |
| AI content can be aesthetically satisfying. | +0.07 |
| AI can never fully replace humans. | +0.13 |

Correlation coefficients were calculated using Spearman’s $\rho$ . Values of $|\rho| \geq 0.10$ indicate a *weak*, and $\geq 0.30$ an *intermediate* relationship.

Participants exhibiting critical or cautious attitudes toward AI tended to show longer and more numerous fixation patterns, whereas those with positive attitudes evaluated the content through shorter, faster, and smoother eye movements. This finding supports the “visual cognitive alignment” hypothesis, which posits that *the more a visual stimulus aligns with an individual’s beliefs, the less cognitive effort is required to interpret it.* Thus, the weak-to-moderate correlations observed between attention metrics and attitudinal scores suggest that perceptual– emotional alignment may function as a guiding mechanism in visual cognition, shaping how individuals process and evaluate AI-generated versus human-made imagery.

## Discussion

This study employed a within-subject experimental design with graphic design students to jointly examine the effects of human-made and AI-generated (artificial intelligence) visuals across three dimensions: attention (eye-tracking), emotionality (facial coding), and subjective evaluation (survey). The findings revealed that human-made visuals elicited longer viewing durations, higher fixation counts, and broader spatial scanning patterns, whereas AI-generated visuals induced more intense but shorter and center-focused attention. In pairwise comparisons, human productions were preferred in terms of aesthetics, originality, and narrative quality, while AI-generated visuals exhibited a clear advantage in salience (attention-grabbing appeal). Facial coding results showed that the mean *k* coefficient fell within a low-to-moderate range; the relatively higher *k* values for AI-generated content indicated sudden and short-lived emotional arousal, whereas human-made visuals elicited lower but more consistent emotional patterns. The following section connects these findings to theoretical and applied literature, discussing the main contributions and points of tension.

Eye-tracking results demonstrate that AI-generated content effectively captures initial attention, yet this attraction does not always translate into sustained exploratory engagement. This distinction supports the argument that generative models, while capable of optimizing formal properties such as high resolution, smoothness, and compositional centrality, may still possess limitations in conveying semantic and intentional layers. Recent advances in visual generation technologies—such as text-to-image diffusion, text-to-3D pipelines, and hybrid rendering systems like Latent Diffusion, NeRF, and Gaussian Splatting—have markedly improved form and visual fidelity (13,44,46,47,63). Nevertheless, ongoing debates persist regarding controllability, scene coherence, and narrative integrity (40,72,73). The broader scan paths and higher fixation frequencies observed for human-made visuals suggest that such works offer aesthetic complexity and traces of human intentionality, prompting viewers to enter a meaning-making phase characterized by longer and more exploratory attention patterns.

Facial coding analyses revealed that AI-generated visuals produced higher but more fluctuating *k* values, whereas human-made visuals resulted in lower yet more stable emotional patterns. This duality implies that AI imagery can evoke sudden arousal through novel or unexpected visual attributes, but such responses may not evolve into sustained emotional resonance. When considered alongside discussions on how social and cultural contexts frame perception (2,8,32) and studies highlighting the conditional trust in automated or algorithmic decision-making (67), the results align with participants’ pragmatic yet cautious stance toward AI. Previous research on student samples has similarly reported an optimistic but careful calibration of the perceived benefit–risk balance regarding AI use (74,75), further reinforcing this pattern of measured acceptance grounded in both aesthetic judgment and technological awareness.

The overall attribution accuracy of approximately 68% indicates that a considerable portion of participants perceived AI-generated images as “human-made.” This finding aligns with ongoing debates in contemporary art and design regarding AI’s growing capacity to imitate human aesthetics (16,18,66,76). It also suggests that the issue of perceptual authenticity is culturally situated: as AI attains greater “human-like visibility,” attribution ambiguity tends to increase (32,68). Given prior evidence that perceptual transparency and intention inference influence performance and acceptance in social and collaborative contexts (35,77), it appears that the acceptance of AI within aesthetic domains is similarly sensitive to context and explainability.

The correlations between Likert-scale attitudes and fixation metrics were weak (|ρ| ≈ .06–.13), yet the observed trends were consistent: critical or human-oriented attitudes were associated with longer and more numerous fixations, whereas pragmatic or instrumental attitudes were linked to shorter and smoother eye movements. This suggests that the relationship between cognitive attention and emotional or ideological stance may not be strictly linear. The partial dissociation between perceptual processing and evaluative judgment is consistent with the literature on human-centered perceptual (78,79). Likewise, studies on perceptions of AI in educational and professional contexts (80–83) similarly report the prevalence of ambivalent attitudes situated within a utility–trust–ethics triad.

Our findings propose the concept of a “salience–meaning gap”, suggesting that AI-generated visuals are exceptionally strong in salience (initial attention capture) but relatively limited in meaning (semantic depth, intentionality, narrative coherence). This proposition implies that while recent advancements in generative modeling (12,13,46,72,84–87) have rapidly elevated surface-level quality and formal precision, they still require greater design governance, interpretability, and interactive control to address the human creativity layers involving expression and intent.

In practical terms, this means that AI holds an advantage in visually attention-driven domains such as advertising and social media, whereas human design—or human–AI co-design— remains essential in areas that demand brand narrative, cultural coding, and emotional engagement (16,31,62,88,89).

In summary, the study demonstrates that AI productions excel in formal visual salience, while human productions maintain strength in semantic and conceptual depth. Emotional responses to AI visuals tend to be intense but fleeting, whereas those to human-made visuals are subtle yet consistent. This duality suggests that AI plays a complementary role within the design ecosystem and that human–AI co-design frameworks, supported by explainable and controllable production strategies, represent a promising approach to bridging the gap between visual salience and meaning in future creative practices.

## Conclusions, Limitations, and Future Directions

This research examined how human-made and AI-generated visuals are evaluated on perceptual, attentional, and emotional levels through a multimodal approach integrating eye-tracking, facial coding, and attitudinal data. Regarding RQ1, the results revealed significant differences in gaze behavior between AI-generated and human-made visuals. Human-created productions elicited longer viewing durations, higher fixation counts, and broader spatial scanning patterns, indicating deeper cognitive engagement and greater aesthetic complexity. Conversely, AI-generated visuals produced shorter but more narrowly focused fixations, suggesting strong perceptual salience yet limited exploratory attention. These findings demonstrate that human-made designs stimulate more extensive meaning-making processes, while AI visuals primarily attract surface-level attention through compositional regularity and technical precision.

In response to RQ2, facial-coding analyses showed that AI-generated visuals elicited stronger but less stable emotional responses (κ = 0.16), whereas human-made visuals generated weaker yet more consistent affective reactions (κ = 0.04). This pattern indicates that algorithmic productions often provoke momentary curiosity or surprise due to their unfamiliar visual structure, yet they fail to sustain long-term emotional resonance or aesthetic immersion. The non-significant correlation between attention and emotion (ρ = −0.015) further suggests that cognitive attention and affective involvement operate as independent but complementary processes within aesthetic perception.

Addressing RQ3, participants’ attitudes and authorship judgments were found to partially modulate both attention and emotional responses. Individuals who held more critical or human-centered views of AI demonstrated longer and more frequent fixation patterns, reflecting analytical and evaluative engagement. Those with more favorable attitudes toward AI exhibited shorter and smoother scanning behaviors, indicating visual fluency aligned with positive expectations. Moreover, the frequent misattribution of AI-generated visuals as human-made (68.4%) supports the existence of a perceptual authenticity bias, in which viewers intuitively associate aesthetic quality with human intention. Taken together, these results show that AI-generated productions excel in perceptual strength but remain limited in emotional depth and narrative coherence. While human-made designs evoke richer semantic processing and sustained engagement, AI imagery though visually powerful functions as an “aesthetic intermediary form” that is technically advanced yet emotionally incomplete. This study thus advances empirical understanding of how attention, emotion, and belief systems jointly shape the perception of algorithmic creativity and highlights the need for future human–AI co-design frameworks capable of bridging the gap between visual salience and aesthetic meaning.

The results of this study indicate that AI-generated productions demonstrate strong aesthetic performance at the perceptual level yet fail to reach the same depth of emotional and narrative engagement as human creativity. While AI productions are visually striking and technically sophisticated, they have not yet achieved the capacity to fully represent human-specific layers of meaning, narrative intentionality, and emotional coherence. Therefore, in their current state, AI productions can be conceptualized as an “aesthetic intermediary form” technically advanced but emotionally incomplete.

This research has several limitations. The participant group consisted exclusively of graphic design students from a single state university, which restricts the generalizability of the findings. Moreover, the limited visual set used in the experiment (two videos, two static images, and three pairwise comparison screens) did not encompass a wide range of styles, contents, or production modes. In some cases, participants’ ability to guess the creator of certain visuals may have introduced label awareness, potentially shaping their perceptual expectations. Furthermore, the facial coding data captured only short-term micro-expressions, which limits the assessment of long-term emotional responses or sustained aesthetic reflections.

Several recommendations can guide future investigations. First, researchers should employ broader and more diversified stimulus sets to examine the aesthetic perception of different types of human and AI-generated productions. Experimental designs in which the same visuals are alternately labeled as “human-made” or “AI-generated” could reveal how beliefs about authorship influence viewer perception. Additionally, studies incorporating repeated or prolonged exposures may help determine whether AI-generated visuals induce perceptual saturation or authenticity fatigue over time. Future research could also combine eye-tracking with other neurophysiological measures such as EEG, pupillometry, or fNIRS to provide deeper insight into the neuroaesthetic underpinnings of attention and affective engagement. Finally, cross-cultural and expertise-based comparative analyses could shed light on how the aesthetics of AI-generated visuals are perceived differently across cultural and experiential contexts.

In conclusion, this study provides consistent evidence that AI-generated productions excel in formal visual salience, whereas human-made productions maintain superiority in semantic and emotional depth. Future research that integrates human–AI co-design frameworks, label manipulation experiments, longitudinal observation, and multimodal neurophysiological analyses can enhance theoretical clarity and provide evidence-based guidance for design practice. Within this framework, AI emerges not as a substitute but as a complementary creative partner that, when combined with human intentionality, can help bridge the gap between attention and meaning in the evolving field of design and visual communication.

## Acknowledgments

The authors gratefully acknowledge the support of the Turkish Cooperation and Coordination Agency (TİKA) and the Artificial Intelligence and Media Laboratory at Ankara Yıldırım Beyazit University, within the scope of the project titled *“Algorithms and Human Communication: New Communication Paradigms with Artificial Intelligence.”*

## Author Contributions

Conceptualization: M.A.

Data curation: K.K.Y., E.N.C.

Formal analysis: K.K.Y., S.K.

Investigation: K.K.Y., M.B.

Methodology: M.A., K.K.Y.

Project administration: M.A.

Resources: M.A., K.K.Y.

Software: K.K.Y., E.N.C.

Supervision: M.A.

Validation: M.A., S.K.

Visualization: E.N.C., K.K.Y.

Writing – original draft: K.K.Y.

Writing – review & editing: M.A., S.K., M.B., E.N.C.

## Availability Statement

Anonymized and derived datasets supporting the findings of this study have been deposited in the Zenodo repository (DOI:10.5281/zenodo.17316598) and are publicly available under the Creative Commons Attribution 4.0 International License (CC BY 4.0). Raw gaze samples and potentially identifiable materials remain protected and are available only upon reasonable request and a data use agreement, in compliance with institutional ethics guidelines.

## Ethics Statement

The study was conducted in accordance with the ethical principles outlined in the Declaration of Helsinki. All participants were informed about the study’s purpose, procedures, confidentiality measures, and their right to withdraw at any time, and each provided written informed consent prior to participation. The research protocol was reviewed and approved by the Scientific Ethics Evaluation Committee of Selçuk University, Faculty of Communication (Decision No. 2025/16-03, dated July 30, 2025).

## Supplementary Information

Supplementary materials related to this study are available in the associated public repository described in the Data Availability section.

